# KSTAR v1.2: A faster and more and accessible KSTAR for kinase activity inference

**DOI:** 10.64898/2026.09.14.751568

**Authors:** Sam Crowl, Joseph-Levi Custer, Gabriela Salazar Lopez, Candace Lei-Dadey, Adrian A. Shimpi, Kristen M. Naegle

**Affiliations:** Biomedical Engineering, University of Virginia, Charlottesville, 22903, Virginia, USA; Department of Genome Sciences, University of Virginia, Charlottesville, 22903, Virginia, USA

**Keywords:** post-translational modifications, database, proteins

## Abstract

**Motivation:** KSTAR is an algorithm with high flexibility for inferring kinase activity from any phosphoproteomic pipeline. However, in its first instantiation (v0.1) it requires Python programming and lots of memory and computational resources.

Hence, we wished to improve speed and accessibility for broader uptake by researchers.

**Results:** Here, we provide an updated algorithm that improves speed and memory, without affecting accuracy, along with some new features for increased usability and insight. KSTAR v1.2 has also been integrated into Galaxy for programming-free activity analysis and ProteomeScout for dataset preparation and interactive plotting.

**Availability and implementation:** **KSTAR is available at** https://github.com/NaegleLab/KSTAR **or on Galaxy on** https://usegalaxy.org/. **KSTAR Network resource assets are managed on Figshare at:** https://doi.org/10.6084/m9.figshare.14944305.

## Introduction

Kinases are key regulatory enzymes that make up a complex network of signaling pathways, allowing cells to quickly adapt to their environment and exhibit precise control on the majority of biological processes. Dysregulation of kinases is a common hallmark of many diseases, including cancer, where a large number of therapeutics targeting kinases have been approved for cancer treatment (7; 1). As such, having tools to effectively measure kinase activity and its regulation are critical for understanding cell behavior and disease, but kinase regulation is often complex, making it difficult to obtain direct measurements of kinase activity. Instead, kinase-substrate based approaches using discovery-based mass-spectrometry datasets have become the preferred approach, with numerous tools like KSEA (2), PTM-SEA (9), and KinaseLibrary (6; 15) being developed to use phosphorylation of a kinase’s substrates as indicators of activity. Most of these approaches either rely on known kinase-substrate interactions (which are sparse and tend to be skewed towards well studied kinases), motif-based predictions (which are missing important context about kinase interactions), or more complex kinase-substrate predictions from tools like NetworKIN (5) (which are plagued by skews towards well studied substrates and kinases (14)). Many of these approaches also rely on relative measurements with fold changes as quantification, making them more difficult to apply to other experimental settings such as patient-specific datasets.

We previously developed KSTAR (Kinase-Substrate Transfer to Activity Relationships) to predict kinase activity from phosphoproteomics data, which can be used for both differential or single sample experiments, and overcomes limitations and bias of kinase-substrate prediction networks (3). KSTAR statistically evaluates the “footprint” of kinase activity by measuring the enrichment of a kinase’s substrates in a dataset. It produces an activity score – the larger the score, the more evidence there is for activity – and a significance based on a measured false positive rate. In addition to its unique flexibility, KSTAR is more robust to study bias and data loss (3) than other algorithms, which comes from two key aspects of the algorithm. First, we are able to use more of a phosphoproteomic dataset by using predicted kinase-substrate networks from NetworKIN (5) instead of annotations, aggregating enrichment across 50 representations of heuristically pruned networks to account for the hypothetical nature of the networks. Second, we measure false positive rates by using decoy experiments that match the size and study bias of the experiment in hand, accounting for experimental specific study bias composition. Based on simple benchmarking (ability to recover expected kinases from perturbation experiments), KSTAR performs similarly for serine/threonine activity compared to other prediction algorithms, but was significantly better for predicting tyrosine kinase activity. Hence, KSTAR provides major advantages in tyrosine kinase activity space, and across the board, advantages in robustness, flexibility, and reduced study bias, compared to other algorithms.

KSTAR v0.1 was available only in Python and required relatively large computational resources, restricting usability for the larger research community. In KSTAR v1.2, we have taken substantial steps to improve KSTAR accessibility, including increasing speed, lowering memory requirements, and integrating KSTAR into non-programmatic resources (Galaxy (12) and ProteomeScout (4), Figure 1A). Here, we benchmark KSTAR v1.2 and provide a comparison to Kinase Library (6; 15), developed since previously published benchmarking, which demonstrates that KSTAR still outperforms available tools, particularly for tyrosine kinase activity prediction.

**Figure 1.**
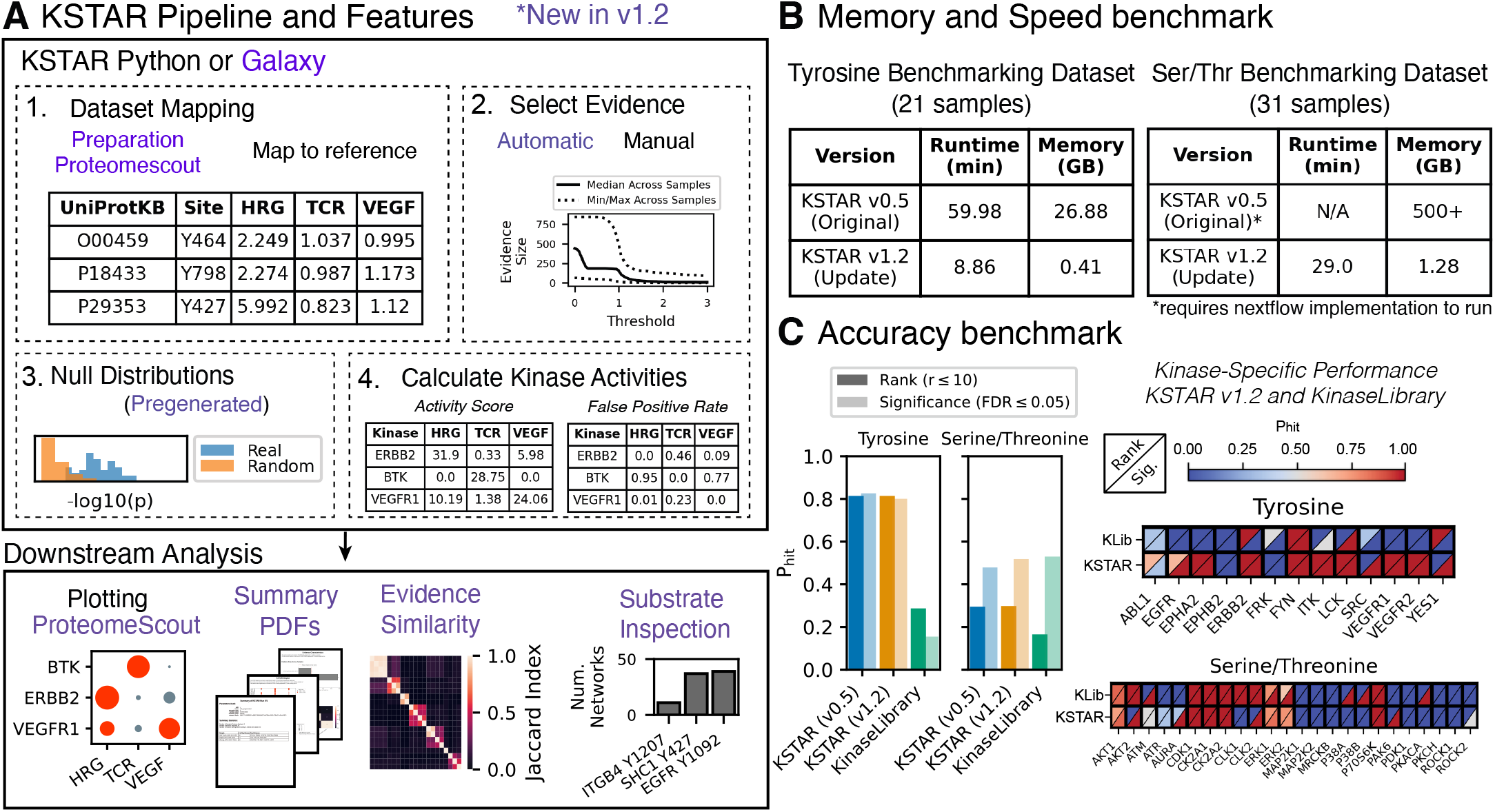
Updates to the KSTAR Pipeline and Performance Assessment. **A)** KSTAR pipeline and downstream analysis options for users – new features denoted in purple. KSTAR generates kinase activity scores and false positive rates based on the statistical enrichment of kinase substrates in a sample relative to what we might expect from a random experiment. The four key steps used in KSTAR to predict kinase activities are indicated. KSTAR-Galaxy provides a programming free environment to run all steps of activity inference and non-programming interaction of plotting has been added to ProteomeScout. Additional new features include: dataset preparation (on ProteomeScout), automatic threshold detection, pregenerated null distributions, summary reports, and evidence inspection. **B)** Runtime and peak memory requirements of KSTAR run on benchmarking dataset for v0.1 and v1.2. Using v0.1, serine/threonine predictions required use of a highly parallel implementation based in nextflow due to significant memory issues. **C)** Ability of KSTAR and KinaseLibrary to recover perturbed kinases from a suite of stimulation and inhibition conditions (21 for tyrosine, 31 for serine/threonine), as described in the original publication (3). Accuracy was defined as *P*_*hit*_ – the fraction of perturbed kinases identified as differentially active, either based on activity rank (in the top 10 kinases) or significance (p or FPR ≤ 0.05). KSTAR v1.2 and KinaseLibrary kinase-specific performance also provided. Performance for each kinase is shown based on Rank and Significance, where *P*_*hit*_ can be a fraction, if more than one benchmark dataset is used to test accuracy for that kinase in the benchmark.

## Methods

### Pregeneration of Random Activity Null Distribution

A principal limitation of the original KSTAR implementation was the time and memory required for large datasets with many samples.

This time and memory issue is predominantly incurred as a result of the ensemble of networks and the customized random null distribution made for each condition in a dataset. For a single sample experiment, KSTAR needed to run a total of 375,000 statistical tests to generate the null distribution for tyrosine kinase activity prediction (150 random experiments x 50 networks x 50 tyrosine kinases). An even greater number of calculations was necessary for the 140 serine/threonine kinases available for KSTAR predictions. To circumvent this issue, we pregenerated multiple sets of random experiments with defined experimental properties, which are matched to the real experiment at runtime (total number of sites and the distribution of study bias). We used benchmarking to evaluate the feasibility of pregeneration and set sizes needed for providing sufficient matches and found the use of pregenerated experiments did not significantly alter activity predictions (see Benchmarking section). Importantly, it reduced the time required to perform predictions by 80%, from 1.7 to 0.4 minutes per sample for tyrosine kinases and 6.8 to 1.4 minutes per sample for serine/threonine kinases. We pregenerated a set of random experiments that match typical experiment sizes and have included these in the KSTAR assets that are installed during KSTAR configuration. For experiments that are beyond the reasonable approximation of either pregenerated sizes or study bias distributions, KSTAR falls back to generating random experiments on the fly, to ensure accuracy, but now includes the option for users to store these locally (i.e. becoming “pregenerated” samples for use in the future).

### KSTAR-Galaxy for a non-programmatic interface

While we have worked to improve the computational load and ease of use of KSTAR as a programmatic tool, its programmatic interface still poses a barrier for researchers not proficient in coding or Python. To enable greater accessibility to KSTAR, we constructed a web-based tool in the Galaxy environment (12). Integration into Galaxy provides several key advantages for applications of KSTAR – 1) many of its tools are already widely adopted by many researchers in both the genomics and proteomics spaces, so many researchers are familiar with the user interface and may be more likely to find and adopt other tools in the ecosystem, and 2) it allows for easy construction of workflows across multiple tools, which will allow users to run Galaxy tools starting from processing of mass spectrometry data files (with tools like MaxQuant (13), FragPipe (8), and EncyclopeDIA (11)) to kinase activity prediction with KSTAR. Importantly, the tool will run KSTAR from start to finish, including properly formatting the input datasets (given the many ways that peptide sequences are annotated), mapping data to the KSTAR reference phosphoproteome, and finally calculating activity prediction. It generates reports that provide researchers with an understanding of possible errors, the size of the evidence used for each experimental column, the overlap of evidence between columns in a dataset, and plots of the full kinase activity matrix. The algorithmic midpoints are available for download from Galaxy, including the mapped experiment, the mapping logs, the binarized experimental assignments (i.e. the evidence included for each data column based on the threshold), and the two files that represent activity scores and false positive rates, which can be immediately used in a web-based plotting tool we integrated into ProteomeScout (4).

### KSTAR streamlining through ProteomeScout: Preparation and Plotting

Galaxy is excellent for running pipelines of analysis, but is not built for real-time user interaction. To improve activity visualization, we created a plotting tool on ProteomeScout, an open source database and interface for post-translational modifications and their experiments (https://proteomescout.research.virginia.edu/kstar/) (10; 4). This tool allows users to upload their KSTAR activity scores, obtained from either the Galaxy tool or Python, and creates live activity plots with high control over sample inclusion and ordering, kinase inclusion and ordering, figure size, and false positive rate settings. The plot can be saved in common formats (such as PNG and SVG). Also on ProteomeScout, we integrated a preparation approach which can map non-UniProt identifiers to UniProt, along with handling various peptide formats and indicators of phosphorylation.

### Thresholding and evidence selection tools

A key aspect of interpreting and using KSTAR activity predictions is dependent on what evidence to use in a dataset for kinase activity calculation. For example, in datasets where there is relative quantification, such as in a perturbation experiment with stimulation or inhibition, selecting a threshold based on that relative quantification (e.g. selecting sites above some threshold value or sites below some threshold value) will change the interpretation of the activity outcomes. Working with scientists across a wide variety of applications, we found a common theme was the desire to find a threshold that highlights differences between conditions in a dataset. We therefore created an evidence diversification approach, which reports the overlap in evidence, based on Jaccard Index, and the size of the evidence for each column across a range of thresholds. Selecting a threshold that maximizes differences, while maintaining sufficient evidence, can then be used in activity analysis. We integrated this threshold selection into both the Python and KSTAR-Galaxy packages as “automatic” thresholding where the algorithm selects the threshold automatically.

## Results

To assess performance of KSTAR v1.2, we utilized the benchmarking dataset previously developed for measuring kinase activity accuracy (3), which consists of kinase perturbation experiments for both tyrosine (21 total conditions) and serine/threonine kinases (31 total conditions). We utilized this dataset to assess improvements to computational load and to ensure KSTAR accuracy was unaffected by memory and speed improvements. We additionally benchmarked a new approach to kinase activity prediction (KinaseLibrary), which was published since the last accuracy benchmark (3).

### Time and memory requirements of KSTAR

One of the key goals of the newest KSTAR changes was to improve the computational load required for KSTAR, especially useful for integration into Galaxy, which we achieved through a combination of the pregeneration of the null model and general code optimization. Using the benchmarking dataset, we compared KSTAR run metrics obtained with the original and optimized implementations of KSTAR (Figure 1B). Using four cores for multiprocessing on a Windows-based Lenovo laptop, the original implementation required 26.8 GB of peak memory and took 59.98 minutes to run on the tyrosine benchmarking dataset (21 conditions). The optimized version of KSTAR, using pregenerated random null distributions, now completes in 8.1 minutes and only requires 0.5 GB of peak memory. These memory and time improvements were even more pronounced for serine/threonine predictions, which were infeasible in Python, due to the high memory costs in the original implementation, but now require approximately 1GB of peak memory. Whereas multiprocessing was a necessity in the original implementation, most machines will now be able to generate KSTAR predictions via the Python package, without significant computational time or memory requirements.

### Evaluating KSTAR v1.2 accuracy

We found high reproducibility of KSTAR results between v0.1 and v1.2, with differences being within the expected range that results from the stochastic nature of the random null distribution (Supplementary Figure 1). Overall, there was no statistical difference in the accuracy of v0.1 and v1.2 KSTAR results (Figure 1C, Supplementary Figure 2). In addition to benchmarking the new KSTAR activity, we also benchmarked recent advances in the field, introduced by the large scale motif profiling of kinases (6; 15), which was integrated into a scoring approach called KinaseLibrary. KSTAR performs similarly in serine/threonine prediction, but outperforms KinaseLibrary for tyrosine kinase activity predictions, consistent with comparison to other algorithms (3) – KSTAR correctly identifies 60-80% of perturbed kinases based on either kinase rank, percentile, or statistical significance, while KinaseLibrary was accurate for less than 25% of kinases assessed regardless of criteria (Figure 1C).

## Conclusion

KSTAR v1.2 is significantly faster, less memory intensive, and without affecting accuracy. Despite new advances in the field, it still performs better for tyrosine kinase activity analysis and provides an approach that can be used on any phosphoproteomics platform, including single sample measurements. Now, with programming free usability, we hope that more users will be able to integrate KSTAR into their research.

## Author contributions statement

L.C., S.C., G.S.L, and C.L. developed code and performed analysis. S.C. and K.M.N wrote the manuscript.

## Acknowledgments

We would like to thank the Galaxy team, especially Dr. Jeremy Goecks and Junhao Qiu for their help in adapting KSTAR for Galaxy. We would also like to thank KSTAR users for their feedback.

