## Supplementary Figures for "KSTAR v1.2: A faster and more and accessible KSTAR for kinase activity inference"

### KSTAR: Advancing the Kinase Activity Field

### List of Figures

**FigureS1** Impact of pregenerated null distributions on final activity scores . . . . . . . . . 2

**A**

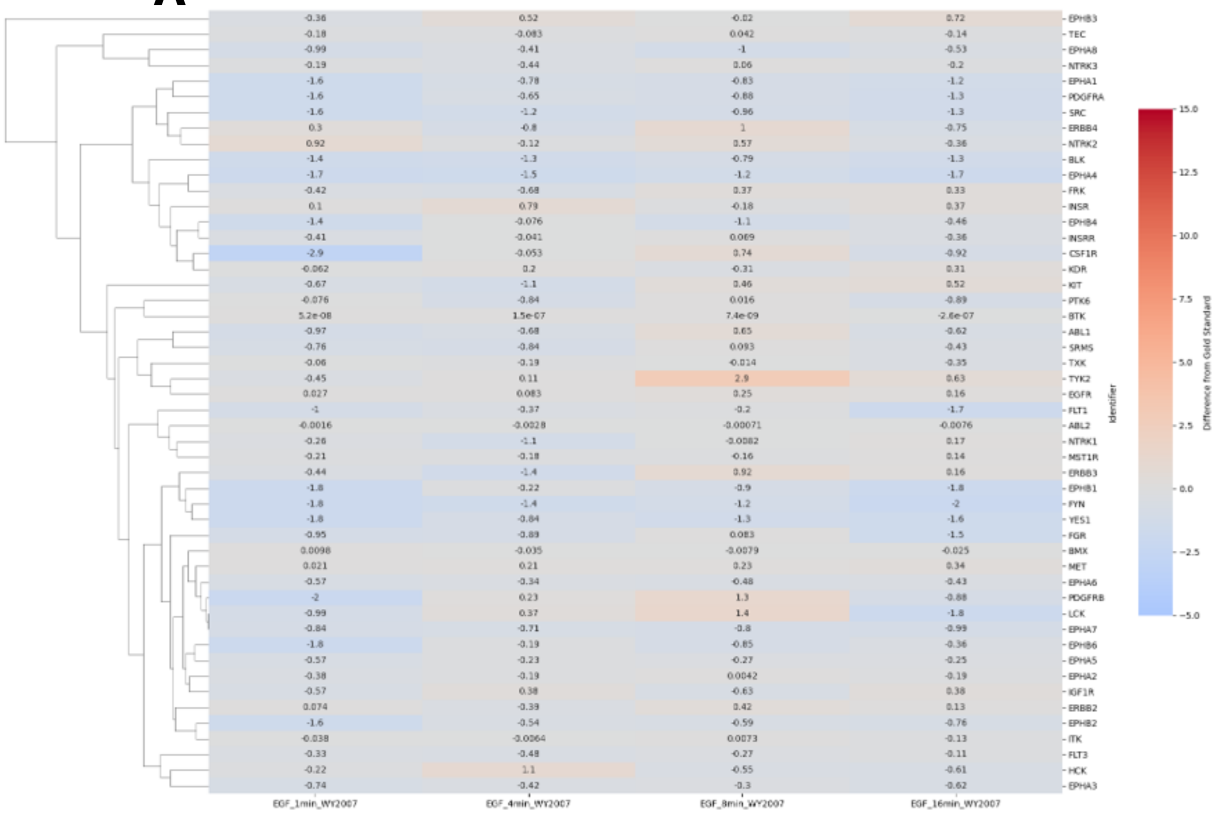


**B**

| Sample Name | Pearson Correlation | P-Value |
|---|---|---|
| EGF_1min_WY2007 | 0.987 | 4.42e-40 |
| EGF_4min_WY2007 | 0.992 | 4.31e-45 |
| EGF_8min_WY2007 | 0.994 | 2.44e-47 |
| EGF_16min_WY2007 | 0.992 | 8.73e-45 |

**Figure S1. Impact of pregenerated null activity distributions on the final activity scores** To ensure that pregeneration of the random activity null distribution did not significantly alter results, we compared tyrosine KSTAR activity scores with and without pregenerated random activities for an EGF stimulation study from Wolf-Yadlin et al. [**?**]. **A)** Differences in KSTAR activity scores between the "gold-standard" (activity scores from original implementation) and the updated KSTAR implementation that uses pregenerated random activity distributions. While there is some variation, these are minor and on par with what we would expect as a result of general stochasticity from generation of the null distribution. **B)** Pearson correlation between activity scores from the original implementation of KSTAR (no pregeneration) and the updated implementation of KSTAR (with pregeneration). Results are in strong agreement.
